# Sulfide toxicity as key control on anaerobic oxidation of methane in eutrophic coastal sediments

**DOI:** 10.1101/2022.02.10.479873

**Authors:** Paula Dalcin Martins, João P. R. C. de Monlevad, Wytze K. Lenstra, Anna J. Wallenius, Maider J. Echeveste Medrano, Martijn Hermans, Caroline P. Slomp, Cornelia U. Welte, Mike S. M. Jetten, Niels A.G.M. van Helmond

## Abstract

Coastal zones account for significant global marine methane emissions to the atmosphere. In coastal ecosystems, the tight balance between microbial methane production and oxidation in sediments prevents most methane from escaping to the water column. Anthropogenic activities, causing eutrophication and bottom water deoxygenation, could disrupt this balance in the microbial methane cycle and lead to increased methane release from coastal sediments. Here, we combined microbiological and biogeochemical analyses of sediments from three sites along a bottom water redox gradient (oxic-hypoxic-euxinic) in the eutrophic Stockholm Archipelago to investigate the impact of anthropogenically-induced redox shifts on microbial methane cycling. At both the hypoxic and euxinic site, sediments displayed a stronger depletion of terminal electron acceptors at depth and a shoaling of the sulfate-methane transition zone in comparison to the oxic site. Porewater methane and sulfide concentrations and potential methane production rates were also higher at the hypoxic and euxinic site. Analyses of metagenome-assembled genomes and 16S rRNA gene profiling indicated that methanogens became more abundant at the hypoxic and euxinic site, while anaerobic methane-oxidizing archaea (ANME), present in low coverage at the oxic site, increased at the hypoxic site but virtually disappeared at the euxinic site. A 98% complete genome of an ANME-2b *Ca*. Methanomarinus archaeon had genes encoding a complete reverse methanogenesis pathway, several multiheme cytochromes, and a sulfite reductase predicted to detoxify sulfite. Based on these results, we infer that sulfide exposure at the euxinic site led to toxicity in ANME, which, despite the abundance of substrates at this site, could no longer thrive. These mechanistic insights imply that the development of euxinia, driven by eutrophication, could disrupt the coastal methane biofilter, leading to increased benthic methane release and potential increased methane emissions from coastal zones to the atmosphere.

## Introduction

Methanogens in marine sediments produce up to 85-300 Tg of the potent greenhouse gas methane per year, which represents 7-25% the of global methane production (Dean et al., 2018; Knittel & Boetius, 2009). However, anaerobic methane-oxidizing (ANME) archaea consume more than 90% of the *in situ* generated methane (Knittel & Boetius, 2009). While coastal zones cover only circa 15% of the total ocean surface area, they account for more than 75% of global marine methane emissions (Bakker et al., 2014; Bange, Bartell, Rapsomanikis, & Andreae, 1994) as an indirect result of high nutrient inputs and burial rates of organic matter (Wallenius, Dalcin Martins, Slomp, & Jetten, 2021). Recent estimates suggest that 5-28□Tg of methane per year is emitted from coastal waters to the atmosphere (Rosentreter et al., 2021). Eutrophication - excess nutrient input - could disrupt the current balance between microbial methane production and consumption. For instance, eutrophication can cause seawater oxygen depletion due to aerobic microbial respiration coupled to degradation of fresh labile organic matter inputs from increased primary productivity, particularly in enclosed basins with a shallow water depth (Middelburg & Levin, 2009). Moreover, eutrophication can lead to shifts in sediment redox conditions and geochemistry, such as vertical compression of the typical marine sedimentary redox-zonation (H.D. Schulz & Zabel, 2006), eventually leading to the release of sulfide into the bottom water (Slomp, 2013) - a condition termed euxinia. Additionally, high inputs of organic matter derived from eutrophication and high sedimentation rates provide increased substrates for methanogenesis (Wallenius et al., 2021). While the combination of these processes is expected to increase methane production in sediments, it remains largely unknown how such processes impact anaerobic methane removal and benthic methane release into the water column. This makes it urgent to mechanistically understand coastal sediment methane dynamics in order to build predictive biogeochemical models of future changes, develop appropriate management strategies, and accelerate the pace of climate action.

ANME archaea are key players in anaerobic methane removal in marine sediment ecosystems ranging from coastal zones to the deep sea (Aromokeye et al., 2020; Susma Bhattarai et al., 2017; Gründger et al., 2019; Pernthaler et al., 2008; Stokke, Roalkvam, Lanzen, Haflidason, & Steen, 2012). Phylogenetically, ANME form three major clades: ANME-1, ANME-2, and ANME-3, recently assigned a putative nomenclature at family and genus level (Chadwick et al., 2022), to which we refer in this section. ANME-1, in the order *Methanophagales*, comprises the family *Methanophagaceae* with at least 6 genera, and are present in a broad range of temperatures, from 2 to 100°C (S. Bhattarai, Cassarini, & Lens, 2019). Members of this clade have been implicated in both methanogenesis and methane oxidation in estuarine sediments (Kevorkian, Callahan, Winstead, & Lloyd, 2021). ANME-2, in the order *Methanosarcinales*, comprises the family *Methanocomedenaceae*, with two genera, *Ca*. Methanocomedens (ANME-2a) and *Ca*. Methanomarinus (ANME-2b), the family *Methanogasteraceae* (ANME-2c), and the family *Methanoperedenaceae* (ANME-2d). A more narrow range of temperatures (4 to 20°C) has been found inhabited by members of this clade (S. Bhattarai et al., 2019). Finally, ANME-3, also in the order *Methanosarcinales*, comprises the family *Methanosarcinaceae* with the genus *Ca*. Methanovorans, and has been reported in colder temperatures (−1 to 17°C) (S. Bhattarai et al., 2019). While ANME-1, ANME-2 and ANME-3 have been implicated in sulfate-dependent anaerobic oxidation of methane (S-AOM) in consortia with a syntrophic sulfate-reducing partner (S. Bhattarai et al., 2019; Boetius et al., 2000), ANME-2d can independently perform nitrate-dependent, iron-dependent and manganese-dependent anaerobic oxidation of methane (N-AOM, Fe-AOM, and Mn-AOM, respectively) (Cai et al., 2018; Ettwig et al., 2016; Haroon et al., 2013; Leu et al., 2020). ANME-2a were enriched in Fe-AOM incubations with sediments of the North Sea (Aromokeye et al., 2020) and their 16S rRNA gene-based abundance correlated to methane and iron concentrations in sediments of the Bothnian Sea (Rasigraf et al., 2019). Moreover, ANME-2a, 2b, 2c, 2d and ANME-3 genomes have genes predicted to encode multiheme *c*-type cytochromes potentially implicated in extracellular electron transfer and iron reduction (Chadwick et al., 2022), suggesting that multiple ANME groups might perform metal-AOM.

The Baltic Sea is highly impacted by eutrophication (Conley et al., 2011; Murray et al., 2019) and has been proposed as a model marine ecosystem indicative of future global changes related to anthropogenic impacts such as oxygen depletion and environmental degradation (Reusch et al., 2018). High methane emissions to the atmosphere from several locations in the Baltic Sea have been documented, in the range of 0.1-3.3 mmol m^-2^ day^-1^, particularly from coastal sites with a shallow sulfate-methane transition zone in the sediment and relatively shallow water depth (Bange et al., 1994; Gülzow, Rehder, Schneider v. Deimling, Seifert, & Tóth, 2013; Humborg et al., 2019). Similarly, significant methane concentrations in the water column (up to 47 nmol L^-1^, (Humborg et al., 2019)), large methane benthic fluxes (up to 2.6 mmol m^-2^ day^-1^, (Sawicka & Brüchert, 2017)), and high porewater concentrations of methane (6 mM, (Sawicka & Brüchert, 2017)) have been reported in the Baltic Sea. Previous studies indicate that ANME are key players in methane cycling in Baltic Sea sediments, with ANME-1 and ANME-2 accounting for S-AOM activity, ANME-2 potentially involved in Fe-AOM, and ANME-1 also implicated in methanogenesis (Beulig, Røy, McGlynn, & Jørgensen, 2019; Iasakov et al., 2022; Meulepas et al., 2009; Rasigraf et al., 2019). However, a mechanistic understanding of environmental and biological factors that impact AOM in the Baltic Sea and other coastal sediment ecosystems remains elusive.

Here we investigated sediments of the eutrophic Stockholm Archipelago (Almroth-Rosell, Edman, Eilola, Meier, & Sahlberg, 2016; Conley et al., 2011; van Helmond et al., 2020), pursuing a better mechanistic understanding of the impacts of differing bottom water redox conditions on microbial methane cycling and associated sediment biogeochemistry. We selected three sites across a gradient of bottom water oxygen concentrations, i.e. oxic: [O_2_]_aq_ > 63 µmol L^-1^, seasonally hypoxic: [O_2_]_aq_ < 63 µmol L^-1^ and euxinic: [O_2_]_aq_ = 0 µmol L^-1^ with free sulfide, for high resolution geochemical characterization and 16S rRNA gene profiling, potential methane production rate measurements, and metagenomic analyses. Our study specifically aimed to identify the main controls on the abundance, distribution and activity of ANME archaea and to elucidate the impacts of differing bottom water redox conditions on methane cycling in these coastal sediments.

## Materials and methods

### Sampling

Sediment cores were collected on board the R/V *Electra* with a Gemini gravity corer (8 cm internal diameter and 1 m length) from June 11 to 13, 2019, at three locations in the Stockholm Archipelago: Site 3 (Sandöfjarden), with oxic bottom waters, Site 5 (Lilla Värtan), with seasonally hypoxic bottom waters, and Site 7 (Skurusundet), with generally euxinic bottom waters (Figure 1). Prior to sediment retrieval, a CTD instrument (SBE 911plus, Sea-bird Scientific, USA) was deployed to determine water column dissolved oxygen concentrations, temperature and salinity. Site coordinates and characteristics are provided in Table 1. At each site, six sediment cores were collected and processed as follows on the same day: one was directly used for microelectrode profiling, one was directly sampled for methane, one was sliced as quickly as possible under a nitrogen atmosphere for porewater and solid-phase characterization, one was sliced under ambient atmospheric conditions (hence oxic) for porosity determination, and two were sliced under a nitrogen atmosphere for microbiological analyses (see detailed methods below).

**Table 1.** Geographical coordinates and characteristics of each sampling site.

| Site | Coordinates (DD°MM'SS'') | Water depth* | Bottom water redox conditions* | Location in the Archipelago** |
| --- | --- | --- | --- | --- |
| **Site 3**<br>(Sandöfjärden) | 59°24'33"N<br>18°31'14"E | 64 m | Oxic | Intermediate |
| **Site 5**<br>(Lilla Värtan) | 59°22'08"N<br>18°05'35"E | 21 m | Seasonally hypoxic | Inner |
| **Site 7**<br>(Skurusundet) | 59°17'55"N<br>18°13'46"E | 27 m | Generally euxinic | Intermediate |

\*Based on monitoring data from the Swedish Meteorological and Hydrological Institute (2019).
\*\*Following a previously described classification system (Almroth-Rosell et al., 2016).

**Figure 1.**
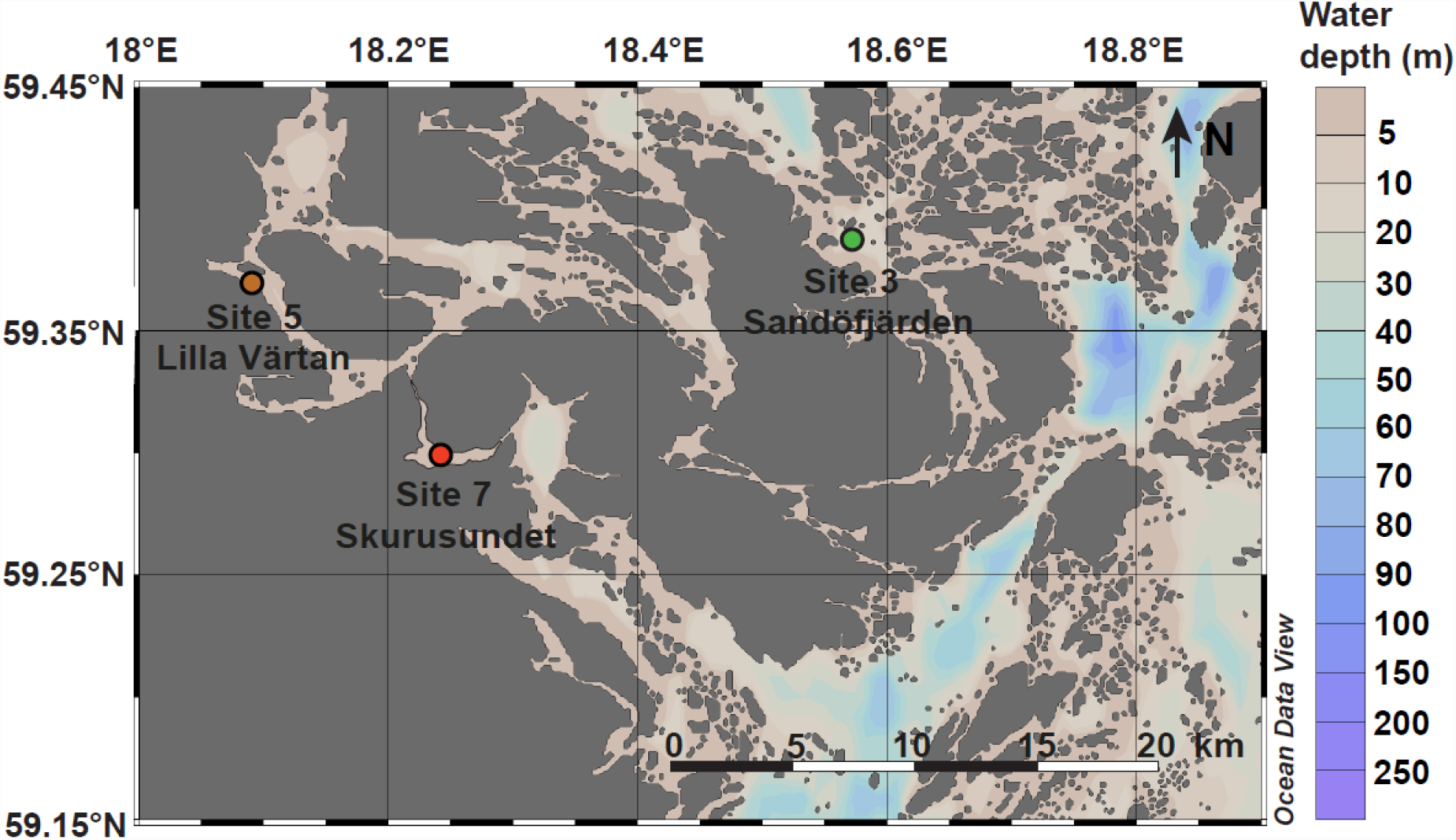
Sampling locations within the Stockholm Archipelago selected for this study. Bottom water redox conditions at each site are color coded as green (oxic), orange (hypoxic) and red (euxinic). This figure was generated using Ocean Data View (Schlitzer, 2018).

### Macrofauna characterization

The top 15 cm of sediment from three cores was sieved over a 0.5□mm mesh size, after which macrofauna was determined at genus level.

### High-resolution depth profiling

High depth resolution (50 µm) profiles of oxygen and pH in the overlying water and surface sediment were obtained from intact sediment cores, using microelectrodes operated by a motorized micromanipulator (Unisense A.S., Denmark). Oxygen was measured immediately, and pH was measured within 1h of core retrieval. For the oxygen microelectrode, a two-point calibration was applied (100% oxygen saturated and nitrogen purged bottom water) using a calibration vessel (Unisense A.S., Denmark, CAL300). For the calibration of the pH microelectrode, three NBS standards (pH 4, 7 and 10) were used. Then, a TRIS buffer was applied for the pH to correct for signal drift induced by salinity effects (A. G. Dickson, Sabine, & Christian, 2007; Andrew G Dickson, 1990).

### Collection of porewater and sediment samples for geochemical analysis

To determine porewater methane (CH_4_) concentrations, samples of the bottom water and sediment were taken with cutoff 10 mL syringes via predrilled holes in the core liner with a depth spacing of 2.5 cm directly after coring. Subsequently, 10 mL of sample was transferred into a 65□mL serum bottle filled with saturated salt solution. Finally, the bottles were stoppered, capped and stored upside down until analysis. For further porewater analysis and collection of anoxic sediments, one core was sliced on board the ship in a N_2_-filled portable glove bag (Sekuroka 940×940×640mm; Glas-Col, USA). Prior to anoxic sectioning of the cores, two samples were taken from the overlying water using a 20 mL syringe equipped with a three-way valve. Subsequently, cores were sliced at a resolution of 0.5 cm (0-2 cm), 1 cm (2-10 cm), 2 cm (10-20 cm), 4 cm (20-40 cm) and 5 cm until the bottom of the core. Sediment was collected in 50 mL centrifuge tubes and centrifuged at 3500 rpm for 20 minutes to extract porewater. Bottom and porewater samples were filtered over 0.45 µm filters in a second N_2_-filled portable glove bag (Sekuroka 690×690×380mm; Glas-Col, USA). Subsamples were taken for (1) sulfide - 0.5 mL of porewater was added to 2 mL 2% zinc acetate and stored at 4°C; (2) dissolved iron (Fe; assumed to be Fe^2+^) and manganese (Mn) - 1 mL of porewater was acidified with 10 μL 30% suprapur HCl and stored at 4°C; (3) sulfate - 0.5 mL sample was collected and stored at 4°C; (4) ammonium - 1 mL sample was collected; and (5) nitrate and nitrite - 1 mL sample was collected, the latter two were stored at -20°C. The residual anoxic sediment after porewater subsampling was stored in N_2_-flushed gas-tight aluminum bags at -20°C until further analysis. Samples from the core that was sliced under ambient atmospheric conditions were dried in an oven (∼1 week at 60°C) to determine the sediment water content. The porosity of the sediment was then calculated assuming a sediment density of 2.65 g cm^−3^.

### Porewater analyses

Methane concentrations were determined with a Thermo Finnigan Trace gas chromatograph (Thermo Fisher Scientific, USA) equipped with a flame ionization detector as previously described (Lenstra et al., 2018). The average analytical uncertainty based on duplicates and triplicates was <5%. Porewater sulfide was determined spectrophotometrically using phenylenediamine and ferric chloride (Cline, 1969). Dissolved Fe and Mn were measured by Inductively Coupled Plasma-Optimal Emission Spectroscopy (ICP-OES) with a Perkin Elmer Avio 500 (Perkin Elmer, USA). Samples were measured with a radial plasma. Argon was used as the plasma gas (10 L min^-1^), nebulizer gas (0.75 L min^-1^) and auxiliary gas (0.2 L min^-1^). Nitrogen was used as the shear gas (25 L min^-1^). The rate of the peristaltic pump was 1.5 mL min^-1^ and the radio frequency generator was set at 1500 watt. Sulfate concentrations were determined by ion chromatography (IC) with a 930 Compact IC Flex (Metrohm, Switzerland), equipped with a Metrosep A Supp 5-150/4.0 Guard column. Blanks and three different QCs were run five times each per series to monitor the detection limit and reproducibility. Average analytical uncertainty based on duplicates was <1%. Ammonium concentrations were determined colorimetrically using indophenol-blue (Solórzano, 1969). Nitrate and nitrite were determined with a Gallery™ Automated Chemistry Analyzer type 861 (Thermo Fisher Scientific, USA) (Doane & Horwáth, 2003) in accordance with NEN-ISO 15923-1 guidelines (https://www.nen.nl/en/nen-iso-15923-1-2013-en-190502).

### Flux estimates

The downward and upward fluxes of sulfate and methane, respectively, into the sulfate-methane transition zone (SMTZ) and the diffusive fluxes of methane across the sediment-water interface were calculated using Fick’s first law of diffusion:

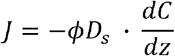

where *j* represents the diffusive flux (mmol m^-2^ d^-1^), *ϕ* represents the sediment porosity, *D_s_* represents the sediment diffusion coefficient for the ambient tortuosity, pressure, temperature and salinity at each site was calculated using the R package *marelac* (Soetaert, Petzoldt, & Meysman, 2010), which implements the constitutive relations previously listed (Boudreau, 1997) and *<u>dC</u>/dz* is: (1) the concentration gradient of sulfate from above the SMTZ into the SMTZ; (2) the concentration gradient of methane into the SMTZ from below and (3) the concentration gradient between the top layer of the sediment and the bottom water. We note that the methane fluxes may be underestimated in sediment sections with high methane concentrations because of loss of methane from the porewater due to degassing during sampling, as observed in previous studies (Egger et al., 2016; Melaniuk, Sztybor, Treude, Sommer, & Rasmussen, 2022).

### Solid-phase analyses

Freeze-dried sediments were ground and homogenized using an agate mortar and pestle inside an argon-filled glovebox and subsequently separated into a fraction that was stored under oxic conditions (oxic fraction) and a fraction that was stored under an argon atmosphere (anoxic fraction). Total organic carbon (TOC) was determined on a subsample of the oxic fraction as previously described (van Helmond, Jilbert, & Slomp, 2018). Briefly, about 200 to 300 mg of sample was decalcified with 1M HCl, after which the residues were dried, powdered and weighed for analysis with a Fisons Instruments NA 1500 NCS analyzer (Carlo Erba, France). TOC was calculated after correction for the weight loss upon decalcification and the salt content of the freeze-dried sediment. Analytical uncertainty, based on duplicates (n=12), was <3%.

The total sedimentary concentrations of Fe and Mn were determined by digestion of ca. 125 mg of the freeze-dried oxic subsample in a mixture of strong acids as previously described (W.K. Lenstra, Klomp, Molema, Behrends, & Slomp, 2021; van Helmond et al., 2020) and analyzed for their elemental composition by ICP-OES. The accuracy (recovery) for Fe and Mn was 91 and 93%, respectively. The average analytical uncertainty based on duplicates (n=14) was <2 % for both Fe and Mn. Two subsamples of the freeze-dried anoxic sample of about 50 mg each were subjected to two different sequential extraction procedures to determine the different solid phase forms of Fe and Mn, respectively.

For Fe, a previously described procedure (Kraal, Dijkstra, Behrends, & Slomp, 2017) was applied, separating sedimentary Fe in: (1) easily reducible Fe-oxides (e.g. ferrihydrite and lepidocrocite), referred to as HCl Fe (III); (2) Fe-carbonates and Fe-monosulfides, (in this setting, with abundant porewater sulfide presumably mainly Fe-monosulfides; (van Helmond et al., 2020), referred to as HCl Fe (II); (3) crystalline Fe-oxides (e.g. goethite and hematite), referred to as citrate-buffered dithionite (CBD) Fe; (4) recalcitrant Fe-oxides, such as magnetite, referred to as Mag Fe; and (5) concentrated HNO_3_-dissolvable fraction (i.e. pyrite), referred to as HNO_3_ Fe. The total Fe concentrations in all extraction steps was determined with the colorimetric phenanthroline method (Saywell & Cunningham, 1937). Average analytical uncertainty for all fractions, based on duplicates (n=12), was <9% for Fe.

For Mn, a previously described procedure (W.K. Lenstra et al., 2021) was applied, separating sedimentary Mn in: (1) poorly ordered Mn-oxides (e.g. birnessite and pyrolusite), referred to as Asc Mn; (2) manganese carbonates, referred to as HCl Mn; (3 & 4) crystalline Mn oxides, referred to as CDB and oxalate Mn; and (5) concentrated HNO_3_-dissolvable fraction, pyrite-bound Mn, referred to as HNO_3_ Mn. The total Mn concentration in extraction steps (1), (2) and (5) was determined by ICP-OES. Steps (3) and (4) were not further analyzed for this study because these fractions generally represented only a minor fraction of the total Mn that was extracted in similar coastal sediments (W.K. Lenstra et al., 2021). Analytical uncertainty, based on duplicates (n=10), was <5% for Mn.

### DNA extractions, amplicon sequencing and 16S rRNA gene analyses

Sediments for DNA sequencing were immediately frozen at -20ºC after on board core slicing under nitrogen atmosphere and were stored at -20ºC for four months until thawing at room temperature for DNA extractions. DNA was extracted from 73 sediment samples retrieved from three cores in total, one core per site, with a depth resolution of 0.5 cm for the top 2 cm, 1 cm until 10 cm depth, 2 cm until 20 cm depth, and 4 cm below that. DNA extractions were performed with the DNeasy Power Soil Kit (Qiagen, Germany) according to the manufacturer’s instructions with two modifications. For the bead-beating step, a TissueLyser LT (Qiagen, Germany) was used for 10 minutes at 50 Hz, and DNA was eluted in 30 µL of autoclaved ultrapure water (Milli-Q Reference Water Purification System, Merck & Co., USA). DNA was quantified with a Qubit 2.0 Fluorometer (Thermo Fisher Scientific, USA). Amplicon sequencing was conducted by Macrogen Europe BV (Amsterdam, Netherlands) on an Illumina MiSeq platform using the MiSeq Reagent Kit v3, producing 2×300bp in paired end readings. For the identification of archaea, the chosen primers were Arch349F (GYGCASCAGKCGMGAAW) and Arch806R (GGACTACVSGGGTATCTAAT) (Takai & Horikoshi, 2000). 16S rRNA gene sequencing data was processed on RStudio v1.3.959 and R v3.6.3 with the following packages: DADA2 (Callahan et al., 2016) v1.9.0, phyloseq (McMurdie & Holmes, 2013) v1.32.0, vegan (Oksanen et al., 2019) v2.5-6, DESeq2 (Love, Huber, & Anders, 2014) v1.28.1, and ggplot2 (Wickham, 2016) v3.3.5. Briefly, primers were removed with cutadapt (Martin, 2011) using the options -g, - G, and --discard-untrimmed. The DADA2 pipeline was then used to trim forward reads at 270 nt and reverse reads at 240 nt based on quality plots. Low quality and contaminant sequences were discarded. After error models were built, sequences were dereplicated and amplicon sequence variants (ASVs) were inferred. Forward and reverse reads were merged and chimeras were discarded. Taxonomy was assigned using the Silva non-redundant train set v138 downloaded from https://zenodo.org/record/3731176#.XoV8D4gzZaQ. ASVs counts and taxonomy tables were then used for data analyses with phyloseq. ASVs were clustered by taxa and relative abundance was calculated and plotted with ggplot2.

### Metagenomic sequencing and data analyses

DNA was extracted and quantified as abovementioned from four homogenized sediment samples from each of the three sites with a depth resolution of 4 cm: 0-4 cm, 9-12 cm, 21-24 cm, and 33-36 cm. These 12 samples were sequenced by Macrogen Europe BV (Amsterdam, Netherlands) using the TruSeq Nano DNA library with an insert size of 350bp on an Illumina NovaSeq6000 platform, producing 2×151bp paired-end reads (5 Gbp per sample). Read quality was assessed with FASTQC v0.11.8 before and after quality-trimming, adapter removal and contaminant filtering, performed with BBDuk (BBTools v38.75). Trimmed reads were co-assembled *de novo* using MEGAHIT v1.2.9 (Li et al., 2016) and mapped to assembled contigs using BBMap (BBTools v38.75). Sequence mapping files were handled and converted using SAMtools v1.10. Contigs at least 1000 bp-long were used for binning with CONCOCT v1.1.0 (Alneberg et al., 2013), MaxBin2 v2.2.7 (Wu, Simmons, & Singer, 2015), and MetaBAT2 v2.15 (Kang et al., 2019). Resulting metagenome-assembled genomes (MAGs) were dereplicated with DAS Tool v1.1.2 (Sieber et al., 2018), manually curated, and taxonomically classified with GTDB-Tk v1.3.0 release 95 (Chaumeil, Mussig, Hugenholtz, & Parks, 2019). MAG completeness and contamination was estimated with CheckM v1.1.2 (Parks, Imelfort, Skennerton, Hugenholtz, & Tyson, 2015).

MAGs were annotated with DRAM v1.0 (Shaffer et al., 2020) with default options, except - min_contig_size 2500 bp for MAGs and 5000 bp for unbinned contigs, and genes of interest were searched in annotation files. Additionally, methyl-coenzyme M reductase alpha subunit-encoding *mcrA* genes were also searched with the HMM PF02745.15 MCR_alpha_N and iron metabolism genes were searched with FeGenie (Garber et al., 2020) v1.0. Only high and medium quality MAGs (>50% complete and less than 10% contaminated) were included in genome-centric analyses, and the entire dataset (binned and unbinned contigs) was considered in gene-centric analyses. For phylogenetic trees, sequences were aligned with muscle v3.8.31 (Edgar, 2004), alignment columns were stripped with trimAl v1.4.rev22 (Capella-Gutierrez, Silla-Martinez, & Gabaldon, 2009) using the option -gappyout, and trees were built with FastTree v2.1.10 (Price, Dehal, & Arkin, 2010) or UBCG v3.0 (Na et al., 2018). Average amino acid identity between selected genomes was calculated using the Kostas Lab tool (http://enve-omics.ce.gatech.edu/g-matrix/index). For this, genomes were gene called with Prodigal v2.6.3 (Hyatt et al., 2010), and amino acid fasta files were used as input.

### Methanogenic incubations

Sediments for incubations to measure potential methane production rates were sliced N_2_-filled portable glove bag and placed into sterile plastic bags (VWR International BV, Amsterdam). The resulting slices had a resolution of 4 cm and were stored anoxically into sealed aluminum bags (Gruber-Folien GmbH & Co. KG, Germany) in the dark at 4ºC for one month from sampling collection until bottles were assembled. For this, 5g of wet sediments were placed in 60 mL-serum bottles, and sulfate-free artificial seawater (ASW) medium pH 7.5 was added to create a 1:1 diluted slurry. The ASW medium was adapted from a previously published (Kester, Duedall, Connors, & Pytkowicz, 1967) to achieve a salinity of 5.3 and contained, per litre, 3.418g NaCl, 1.54g MgCl_2_.6H_2_O, 0.097g KCl, 0.21g CaCl_2_.2H2O, 0.014g KBr, 0.0037 H_3_BO_3_, 0.003g SrCl_2_.6H_2_O. 0.0004g NaF, and 0.028g NaHCO_3_. No trace elements or vitamins were added. From each site, six depths were incubated in triplicate, resulting in 54 serum bottles. Methane production was monitored via injection of 100 µl-headspace samples into an HP 5890 gas chromatograph equipped with a Porapak Q column (80/100 mesh) and flame ionization detector (Hewlett-Packard, USA) with a detection limit of <1 ppm. Each gas sample was measured in triplicate and areas were averaged.

Methane concentrations in the headspace were derived from a calibration curve (R^2^>0.99). The percent of methane in the headspace was converted to molar concentration using the ideal gas law, where one mole of an ideal gas has a volume of ∼22.4 L at room temperature and 1 atm. To calculate the volume (ml) of the liquid in incubation bottles, the amount of added medium (5 mL) was summed to sediment porewater volumes (calculated from dry weight measurements). Dissolved methane in the liquid was then estimated based on Bunsen solubility coefficients extrapolated from experimental data (Yamamoto, Alcauskas, & Crozier, 1976). For 4°C and a salinity of 5.3, the dimensionless solubility was estimated to be 0.0493, resulting in a Henry’s law coefficient (Hcp) of 0.0022 mol L^-1^ atm^-1^ as calculated with equation 12 as previously described (Sander, 2015). From this, we used Henry’s law (C_gas_ = Hcp x P_gas_) to estimate that 1 atm 100% methane at 4°C has a solubility of 2.2 mM in seawater with a salinity of 5.3, and then liquid-dissolved methane concentrations were proportionally adjusted to gas-phase methane concentrations. Gas-phase and liquid-phase methane amounts were summed to obtain the total amount of methane in each bottle for each time point. Potential rates of methane production were calculated using linear regression of methane measurements obtained in the first 8-9 days of sediment incubation. All calculations and measurements are provided as Supplemental Information.

## Data availability

Adapter-trimmed 16S rRNA gene reads and MAGs have been deposited on NCBI under BioProject PRJNA805085. Geochemical data used to generate the figures in this study are provided as Supplemental Information.

## Results

Bottom water redox conditions at the time of sampling were in line with expectations based on long term monitoring data (Table 1; SMHI, 2019), i.e. oxic, hypoxic and euxinic conditions prevailed at Sites 3, 5 and 7, respectively (Table 2). Oxygen penetration into the sediment followed the trend in ambient bottom water redox conditions, with the deepest oxygen penetration at Site 3 and no oxygen penetration at Site 7. Macrofauna (polychaetes and bivalves) were only present at Site 3. While bottom water pH, salinity and temperature were similar across sites, the different bottom water redox conditions were reflected in full profile-averaged sedimentary TOC contents, with the highest TOC observed at the euxinic site, Site 7 (Table 2 and Supporting Information).

**Table 2.** Characterization of sampling sites. BW, bottom water; TOC, total organic carbon.

| Site | Oxygen (µmol L<sup>-1</sup>) and pH at sediment-water interface* | Oxygen penetration depth (mm)* | BW salinity | BW temperature (°C) | Macrofauna | Average TOC (%)* |
| --- | --- | --- | --- | --- | --- | --- |
| **Site 3**<br>(Sandöfjorden) | 150<br>7.4 | 5.3 | 5.8 | 2.1 | *Marenzelleria*<br>sp. and *M. balthica* | 5.1 |
| **Site 5**<br>(Lilla Värtan) | 55<br>7.3 | 1.4 | 4.9 | 2.3 | none | 5.9 |
| **Site 7**<br>(Skurusundet) | 0**<br>7.1 | 0 | 5.3 | 2.8 | none | 8.5 |

\*Full profiles in Supplemental Information.
\*\*At Site 7, sulfide (10 µmol L<sup>-1</sup>) was present in the bottom water.

Porewater sulfate decreased with depth at all sites (Fig. 2). At Site 3, sulfate was nearly completely removed at ∼30 centimeters below the seafloor (cmbsf), whereas at Sites 5 and 7 this removal occurred around 10 cmbsf (Fig. 2). Methane concentrations increased slightly with depth to a maximum of ca. 100 µmol L^-1^ at Site 3. At Sites 5 and 7, in contrast, concentrations of methane increased strongly with depth, reaching values of ∼2 mmol L^-1^. Porewater sulfide was only present at low concentrations (<200 µmol L^-1^) and in a confined zone (10-30 cmbsf) at Site 3. At Sites 5 and 7, however, sulfide concentrations increased rapidly with depth, with the strongest increase and highest concentrations (>1 mmol L^-1^) at Site 7. At Sites 3 and 5, a peak in porewater Fe was observed directly below the sediment-water interface, with the broadest peak and highest maximum values at Site 3 (∼100 µmol L^-1^), followed by a rapid decrease to values around zero. At Site 3, dissolved Fe concentrations increased again when sulfide was depleted at depth. Dissolved Fe was nearly absent at Site 7. A peak in dissolved Mn was observed near the sediment-water interface at all three sites, with maximum concentrations decreasing with increasingly reducing sediments from Site 3 to Site 5 to Site 7. At Site 3, dissolved Mn in the porewater increased again after a subsurface minimum, reaching maximum values (∼170 µmol L^-1^) at depth. At Sites 5 and 7, dissolved Mn was depleted below 10 cm depth. Ammonium increased with depth at all sites, with the highest concentrations (up to 2.5 mmol L^-1^) and most rapid increase at Sites 5 and 7. NO_x_ concentrations ranged from 2.5 to 12.5 µmol L^-1^ in the bottom waters and decreased rapidly with depth in the sediment at all sites.

**Figure 2.**
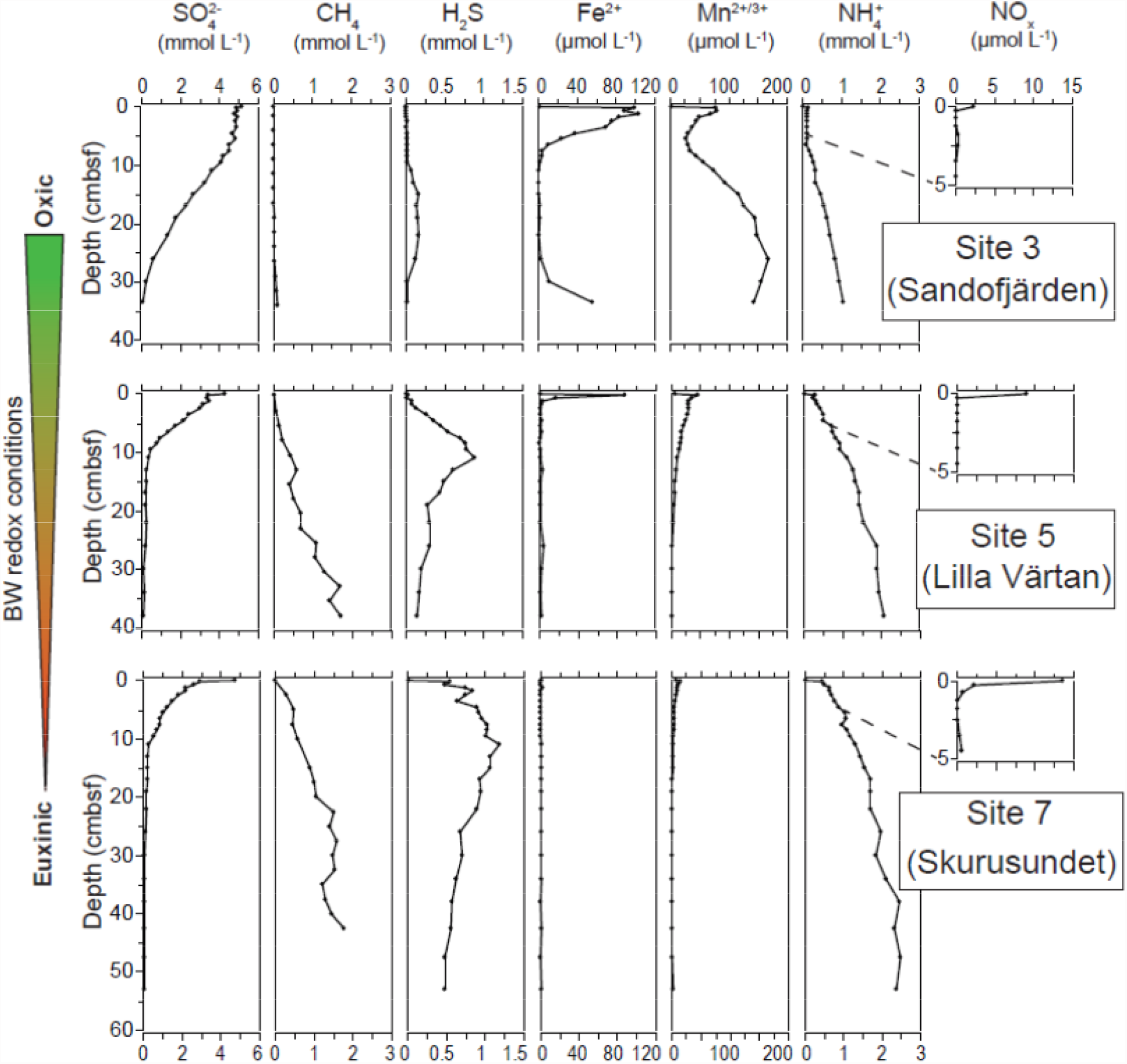
Porewater depth profiles of sulfate (SO_4_ ^2−^), methane (CH_4_), sulfide (H^2^S), dissolved iron (Fe^2+^) and manganese (Mn^2+/3+^), ammonium (NH_4_^+^) and the sum of nitrate and nitrite (NO_x_) at our study sites in the Stockholm Archipelago: Site 3 (Sandofjärden), Site 5 (Lilla Värtan) and Site 7 (Skurusundet). The arrow to the left indicates decreasing bottom water (BW) oxygen concentrations from Site 3 to 7. Cmbsf; centimeters below the seafloor.

The calculated downward flux of sulfate into the SMTZ (Table 3) is highest at Sites 5 and 7 (1.5 and 1.3 mmol m^2^ d^-1^), but still substantial (almost 1 mmol m^2^ d^-1^) at Site 3. The calculated upward flux of methane into the SMTZ at Site 3 is nearly absent, and around 0.5 mmol m^2^ d^-1^ at Sites 5 and 7, which should be regarded as an absolute minimum estimate due to degassing of methane during sampling (Egger et al., 2016; Melaniuk et al., 2022). There is no benthic methane efflux at Site 3. The benthic methane efflux is about seven times larger at Site 7 relative to Site 5 (∼1 vs. 0.15 mmol m^2^ d^-1^, respectively).

**Table 3.** Diffusive fluxes of sulfate and methane (mmol m^-2^ day^-1^) as calculated from porewater profiles (intervals in parentheses), and, for the benthic flux, from porewater and bottom water concentrations (see text). SMTZ; sulfate-methane transition zone.

| Site | Downward sulfate flux into SMTZ | Upward methane flux into SMTZ* | Benthic methane efflux |
| --- | --- | --- | --- |
| **Site 3**<br>(Sandöfjärden) | 0.88 (9.5-19 cm) | 0.08 (29-34 cm) | 0 |
| **Site 5**<br>(Lilla Värtan) | 1.49 (1.75-7.5 cm) | 0.54 (8-13 cm) | 0.15 |
| **Site 7**<br>(Skurusundet) | 1.27 (2.5-4.5 cm) | 0.50 (0-15 cm) | 1.02 |

\*Methane fluxes into the SMTZ at Sites 5 and 7 are likely underestimated because of degassing during sampling.

The sequential extractions dissolved between 70% and 80% of the total sediment Fe at the three sites (Fig. 3). Except for the surface sediment at Site 3, the easily reducible Fe-oxide content of the sediment (HCl Fe(III)) was generally low. Crystalline Fe-oxides (CDB-Fe) accounted for a substantial fraction of the extractable Fe at all sites (15-30%). While Fe-monosulfides (HCl Fe(II)) were the dominant Fe fraction at depth in the sediment at Site 3, both Fe-monosulfides and pyrite (HNO_3_ Fe) accounted for the majority of the extractable Fe at depth in the sediment at Sites 5 and 7.

**Figure 3.**
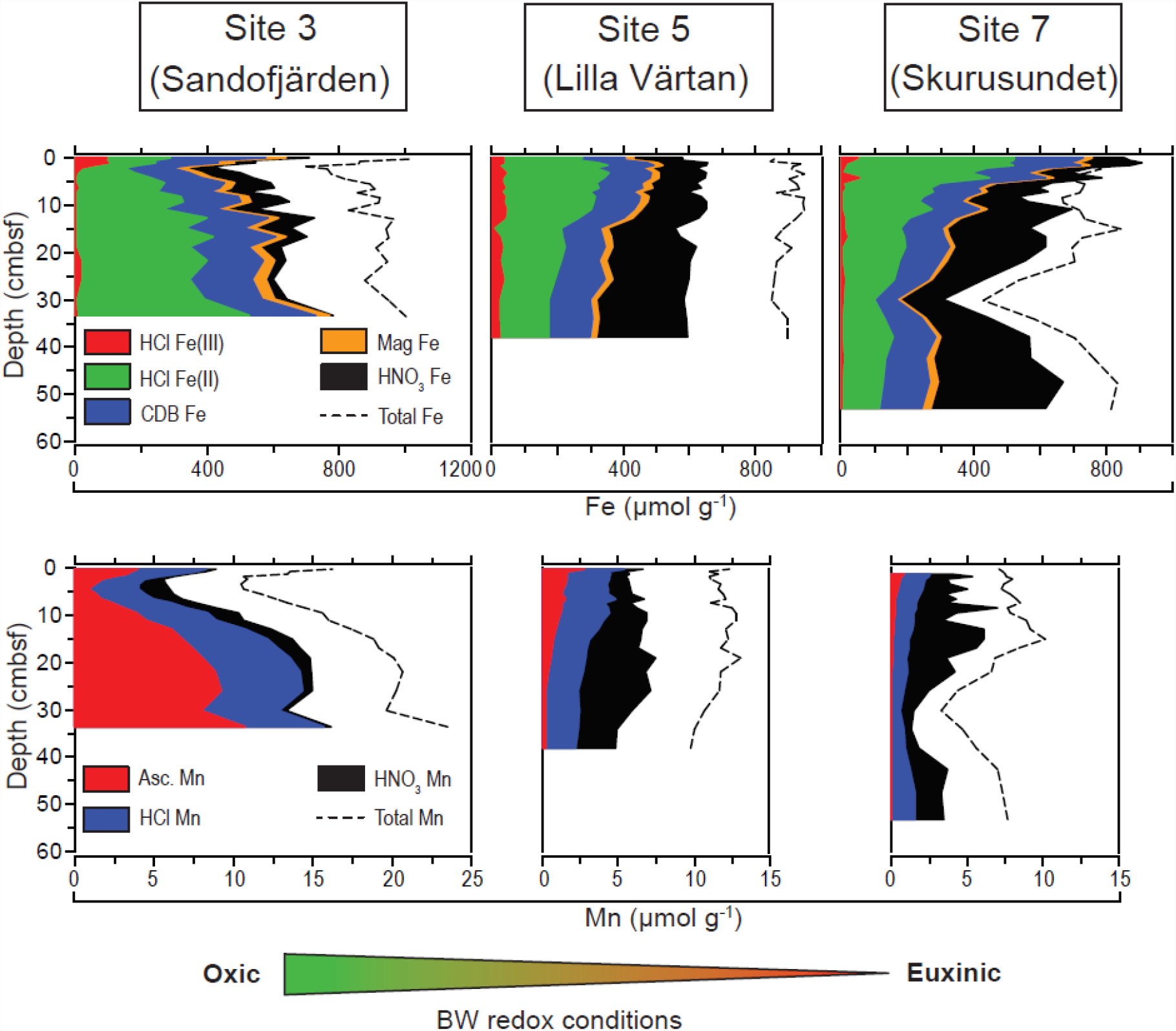
Solid-phase iron and manganese speciation (μmol g^-1^ dry sediment) depth profiles for the three study sites. Easily reducible Fe-oxides are referred to as HCl Fe(III). Fe-carbonates and Fe-monosulfides, but in this setting presumably mainly Fe-monosulfides (van Helmond et al., 2020), are referred to as HCl Fe(II). Crystalline Fe-oxides are referred to as CDB Fe. Recalcitrant Fe-oxides are referred to as Mag Fe. Concentrated HNO_3_-extracted Fe, i.e. pyrite, is referred to as HNO_3_ Fe. Poorly ordered Mn-oxides are referred to as Asc Mn. Manganese carbonates are referred to as HCl Mn. Concentrated HNO_3_-extracted crystalline Mn-oxides and pyrite-bound Mn are referred to as HNO_3_ Mn. The arrow at the bottom indicates decreasing bottom water (BW) oxygen concentrations from Site 3 to 7. cmbsf; centimeters below the seafloor.

About 50 to 60% of the total sedimentary Mn was extracted in the analyzed steps, i.e. steps 1, 2 and 5 (see section on solid-phase analysis), at all three sites. At Site 3, poorly ordered Mn-oxides (Asc Mn) dominated the extracted Mn pool with only a minor (<10%) contribution of pyrite-bound Mn (HNO_3_ Mn). At Site 5, Mn-oxides were never dominant and concentrations were low at depth. Sedimentary Mn could generally be divided into two relatively equal fractions of Mn-carbonate (HCl Mn) and pyrite-bound Mn fractions. At Site 7, pyrite-bound Mn dominated the extracted Mn pool with only a minor (<10%) contribution of poorly ordered Mn-oxides.

Based on the porewater profiles, metabolic potential for anaerobic methane production and consumption was expected. Sites 5 and 7 showed particularly high methane production rates (up 2.3±0.3 and 3.3±0.4 µmol methane g^-1^ d^-1^ respectively at 2 cm depth). Contrastingly, at Site 3, methane production rates did not exceed 0.75±1.3 µmol methane g^-1^ d^-1^ at 18 cm (Figure 4). To underpin the microbial diversity and metabolic potential, sediments were subjected to DNA extraction and high-resolution 16S rRNA gene sequencing, with selected samples also used for metagenomic sequencing. Archaeal 16S rRNA gene sequences were used to generate amplicon sequencing variants (ASVs), which were clustered at family level for relative abundance visualization (Figure 4). *Methanoregulaceae* and *Methanosaetaceae* represented the two most abundant putative methanogenic families. While *Methanoregulaceae* had highest relative abundances of 16% in Site 3 at 34 cm, 38% in Site 5 at 50 cm, and 34% in Site 7 at 30 cm, *Methanosaetaceae* reached 11% in Site 3 at 19 cm, 20% in Site 5 at 42 cm, and 24% in Site 7 at 38 cm. Other identified putative methanogenic families within *Euryarchaeota* included *Methanosarcinaceae, Methanofastidiosaceae, Methanomicrobiaceae, Methanospirillaceae, Methanobacteriaceae, Methanocorpusculaceae*, poorly resolved *Methanocellales* and *Methanomicrobiales* families, *Methanocellaceae*, and *Methanothermobacteriaceae*, and putative methanotroph family *Methanoperedenaceae*. Within the phylum *Verstraetearchaeota*, the putative methanogenic families *Methanomethyliaceae* and *Methanomethylophilaceae* were identified, and, within the phylum *Thermoplasmatota, Methanomassiliicoccaceae*. An ANME-2a-2b family had highest relative abundances, among archaeal sequences, of 32% at 2.5 cm depth at Site 3, 12% at 26 cm in Site 5, and 1% at 50 cm depth in Site 7. The putative ammonium-oxidizing *Crenarchaeota* family *Nitrosopumilaceae* reached 70% relative abundance in Site 3 at 1.25 cm and was minor in the other two sites.

**Figure 4.**
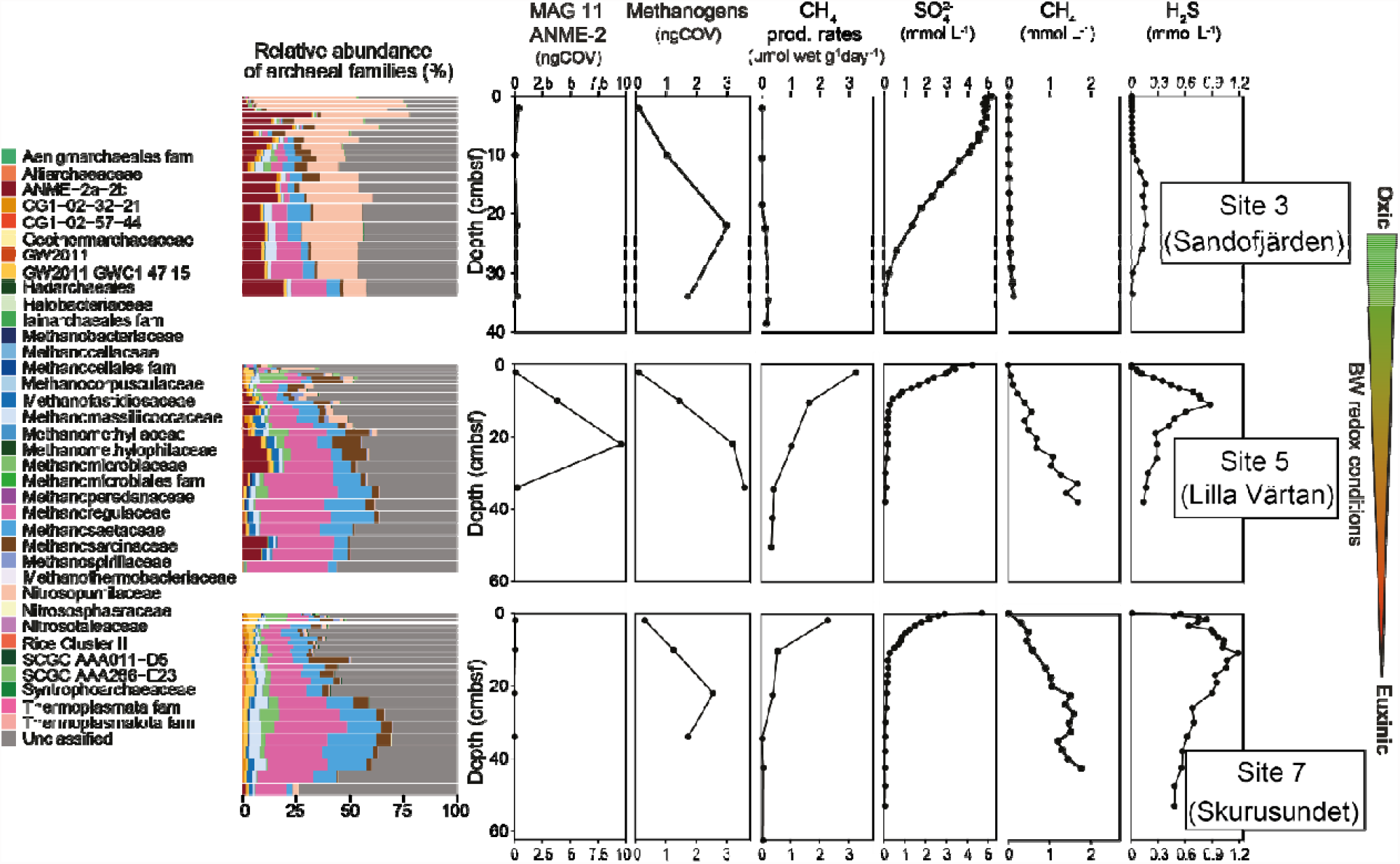
Abundance, distribution and activity of key microbial groups in sediments from the three study sites presented with selected geochemical data. Relative abundances (%) of archaeal families, based on 16S rRNA gene sequencing, are color coded according to the legend to the left and match depths as in other panels. The designation “fam” indicates a poorly resolved family. The thickness of bars corresponds to the depth resolution indicated in materials and methods. The normalized genome coverage (ngCOV) of metagenome-assembled genomes (MAGs) representing methanotrophs (one genome) and methanogens (four genomes) is displayed next, followed by methane production rates in µmol methane wet sediment g^-1^ d^-1^ as measured in triplicate methanogenic incubations, in which error bars are generally smaller than black circles. Sulfate and methane porewater concentrations are expressed in mmol L^-1^. The arrow to the right indicates decreasing bottom water (BW) oxygen concentrations from Site 3 to 7. cmbsf; centimeters below the seafloor.

Metagenome-assembled genomes (MAGs) reconstructed from 12 co-assembled samples were screened for methane metabolism marker genes. No particulate or soluble methane monooxygenase-encoding genes, makers for aerobic methane oxidation, were found in binned and unbinned contigs. Genes encoding a methyl-coenzyme M reductase alpha subunit (*mcrA*), indicative of methane production, were identified in the following four genomes. MAG 009 Methanoregulaceae had potential for hydrogenotrophic methanogenesis, and MAG 010 Methanosarcinaceae had potential for methanogenesis from H_2_ and CO_2_, formate, acetate (*acss*), H_2_ and methanol (*mtaA*), H_2_ and mono-(*mtmBC*) and trimethylamine (*mttC*), but contained two *mcrA* genes. While MAG 015 Methanomassiliicoccales had potential for methanogenesis from H_2_ and methanol (*mtaA*), MAG 016 Methanomassiliicoccales had potential for methanogenesis from H_2_ and methanol (*mtaA*), di-(*mtbC*) and trimethylamine (*mttC*). We also identified an unbinned *mcrA* sequence with a best blastp hit to *Candidatus* Methanofastidiosum methylthiophilus (KYC53403.1 NCBI accession number), which has been proposed to perform methanogenesis from methanethiol, dimethylsulfide, 3-methylmercaptopropionate, and 3-mercaptopropionate (Nobu, Narihiro, Kuroda, Mei, & Liu, 2016). Normalized mapped reads (NMR) values (a proxy for abundance) of *mcrA* genes indicated that MAG 010 Methanosarcinaceae, MAG 015 Methanomassiliicoccales, and MAG 009 Methanoregulaceae accounted for the most significant methanogen *mcrA* NMR changes across sites (Supplemental Figure 1), with highest NMR values in Site 5, then Site 7, and lowest in Site 3. The summed normalized genome coverage (ngCOV) of four methanogen MAGs were highest at Site 5 (3.6x at 34 cm), followed by Site 3 (3x at 22 cm) and Site 7 (2.6x at 22 cm, Figure 4). Similarly, MAG 11 ANME-2 had highest ngCOV at Site 5 (9.5x at 22 cm) below the sulfate-methane transition zone (SMTZ; 0-10 cm) (Figure 4). This MAG ngCOV reached only 0.3x at Site 3 at 2 cm and 0.06x at Site 7 at 2 cm.

Based on phylogenetic analyses, MAG 011, affiliated to the archaeal order *Methanosarcinales*, was identified as representative of an ANME archaeon (Supplemental Figure 2). This genome had 66% average amino acid identity to genome MZXQ01 (NCBI accession number), which represents an ANME-2b archaeon obtained from sediment of the Hydrate Ridge North methane seep (Yu et al., 2018) recently classified as *Ca*. Methanomarinus sp. nov. 1 (Chadwick et al., 2022) (Supplemental Figure 3). MAG 011 was 98.4% complete and 2% contaminated and had potential for a full reverse methanogenesis pathway, as well as acetate production or assimilation via an acetyl-CoA synthase (*acss*) gene (Figure 5). MAG 011 was further analyzed in detail to elucidate its metabolic potential. Genes encoding proteins involved in the reverse methanogenesis pathway were mostly present as single copy, with a few exceptions: (i) *hdrABC* were present in three copies, (ii) a second copy of *mtrAH* was downstream of *mtrX* while *mtrEDCBAFGH* subunits were present in a separate contig, (iii) three copies of the gene encoding the formylmethanofuran-tetrahydromethanopterin N-formyltransferase (*ftr*) were present, and (iv) both molybdenum- and tungsten-dependent formylmethanofuran dehydrogenases were present (*fmdCABDE* and *fwdGBDC*), with *fwdC* present as a separate single subunit and also as a two-copy *fwdC* gene fusion (Supplemental Table 2).

**Figure 5.**
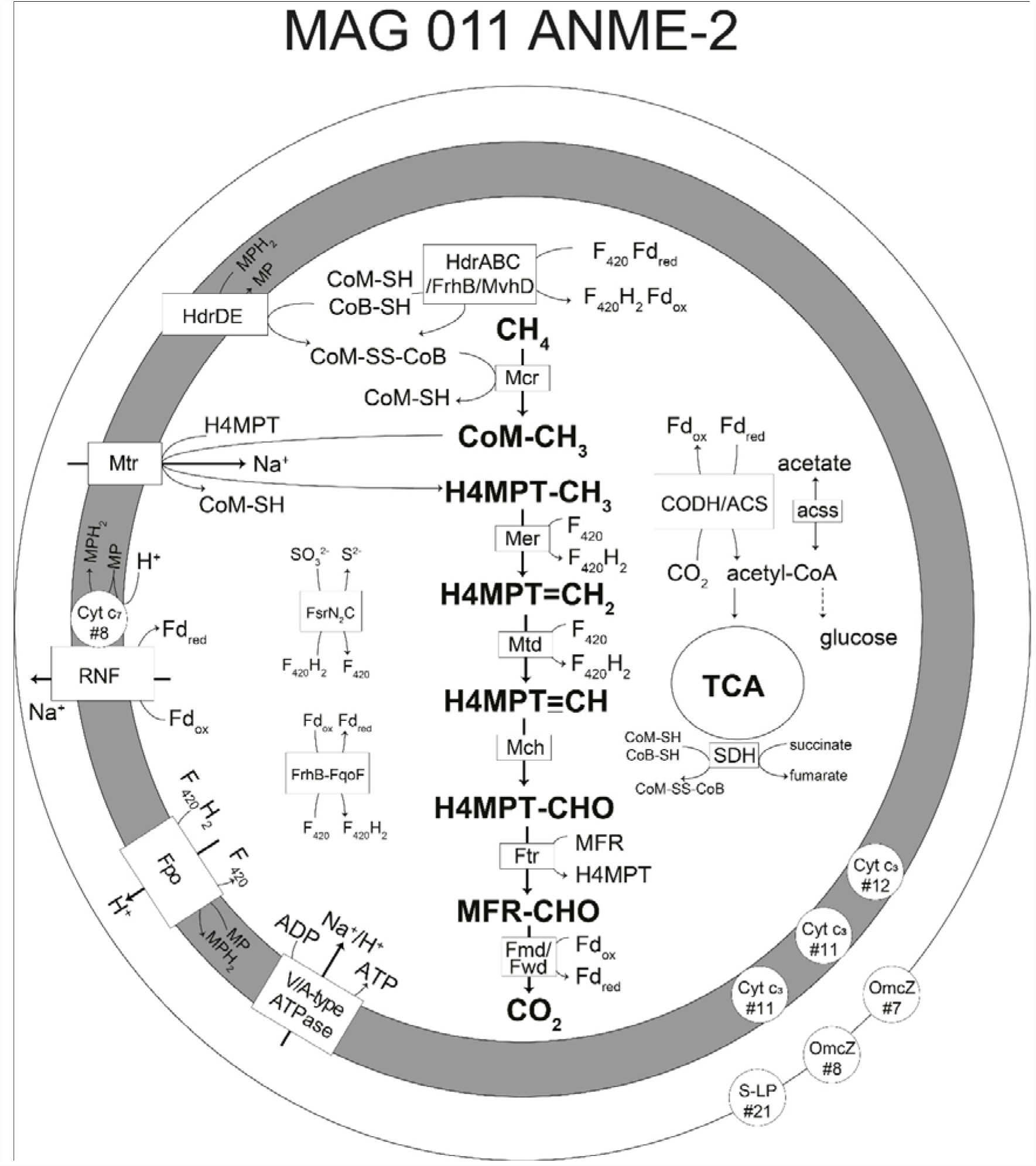
Metabolic reconstruction of MAG 011 ANME-2 based on loci specified in Supplemental Table 2. The grey area represents the pseudo-periplasm, and the outermost circle represents the S-layer. Numbers that follow # indicate the number of heme-binding motifs. Abbreviations are as follows: F_420_, coenzyme F_420_; Fd, ferredoxin; CoM, coenzyme M; CoB, coenzyme B; MP, methanophenazine; HdrABC, cytoplasmic heterodisulfide reductase; Frh, F_420_-reducing hydrogenase; MvhD, methyl viologen-reducing hydrogenase subunit D; HdrDE, periplasmic heterodisulfide reductase; Mcr, methyl-coenzyme M reductase; H_4_MPT, tetrahydromethanopterin, Mtr, formylmethanofuran-H_4_MPT N-formyltransferase; Mer, F_420_-dependent methylene-H_4_MPT reductase; Mtd, F_420_-dependent methylene H_4_MPT dehydrogenase; Mch, Methenyl-H_4_MPT cyclohydrolase; Ftr, Formylmethanofuran: H_4_MPT formyltransferase; Fmd, molybdenum-dependent formylmethanofuran dehydrogenase; Fwd, tungsten-dependent formylmethanofuran dehydrogenase; Cyt, cytochrome; RNF, *Rhodobacter* nitrogen fixation complex; Fpo, F_420_:methanophenazine oxidoreductase; S-LP, S-layer protein; OmcZ, outer membrane cytochrome; FsrN_2_C, F_420_-dependent sulfite reductase; FrhB-FqoF, hypothetical F_420_- and Fd-oxidizing electron-confurcating hydrogenase; CODH/ACS, carbon monoxide dehydrogenase/acetyl-CoA synthase complex EC: 1.2.7.4/ 2.3.1.169); acss, acetyl-CoA synthetase (EC: 6.2.1.1); TCA, tricarboxylic acid cycle; SDH, succinate dehydrogenase.

Nine candidate ferredoxin genes were identified, as well as several FrhB/FdhB/FpoF paralogs. A single subunit *fpoF* could be part of the F_420_H_2_ dehydrogenase complex *fpoDCBAONMLKJ1J2IH*, and an *frhB*-*fqoF* (K00441, K22162) gene fusion could encode a protein to couple ferredoxin oxidation to F_420_H_2_ production. Moreover, three other genes with homology to FrhB of *Candidatus* Methanoperedens nitroreducens strain BLZ1 (Arshad et al., 2015) were found: (i) the first as a single subunit, (ii) the second immediately upstream of *mvhD, hdrA2*, another *mvhD*, then *hdrABCC*, and (iii) the third as 2x*frhB*-*fsrC* fusion (K00441-K00441-K21816). The genes *fsrNC* encode an F_420_-dependent sulfite reductase in *Methanocaldococcus jannaschii* (Johnson & Mukhopadhyay, 2005) (EC: 1.8.98.3), which detoxifies sulfite while reducing it to sulfide. Furthermore, the succinate dehydrogenase membrane subunits *sdhCD* were absent, while *sdhAB* were present and *sdhB* was fused with *tfrB* (K00239, K00240-K18210), which encodes the CoM/CoB-fumarate reductase subunit B (EC: 1.3.4.1) characterized in *Methanobacterium thermoautotrophicum* strain Marburg (Heim, Kunkel, Thauer, & Hedderich, 1998). In *M. thermoautotrophicum*, TfrA harbors FAD-binding motifs and the catalytic site for fumarate reduction, while TfrB harbors one [2Fe-2S] cluster, two [4Fe-4S] clusters, and the catalytic site for CoM-S-H and CoB-S-H oxidation. Therefore, we hypothesize that in MAG 011 ANME-2 electrons from succinate oxidation could be used to generate heterodisulfide for the first step in methane oxidation instead of flowing to the electron transport chain. Finally, a complete Rnf complex was identified, as well as a downstream c_7_ family-octaheme cytochrome as previously reported in ANME-2 (Wang et al., 2014). Other cytochromes were also identified in this genome: three c_3_-family cytochromes containing 11 or 12 heme-binding motifs, an S-layer protein containing 21 heme-binding motifs, and two FeGenie-identified outer membrane cytochrome *omcZ* sequences with 7 and 8 heme-binding motifs, which could mediate extracellular electron transfer to a syntrophic partner or metallic terminal electron acceptor.

## Discussion

All three sites are characterized by a shallow SMTZ, as also observed at other sites in the Stockholm Archipelago (Sawicka & Brüchert, 2017; van Helmond et al., 2020), which is a common feature of depositional coastal sediments (Wallenius et al., 2021). The shallow SMTZ is attributed to a combination of both low salinity, hence low sulfate concentrations, and high rates of organic matter deposition and degradation, culminating in a vertical compression of the redox-zonation. The deeper SMTZ at Site 3, when compared to the other two sites, can be explained by its ambient redox conditions. Oxygen is available in the bottom waters throughout the year, leading to aerobic degradation of organic matter in the surface sediments, whereas anaerobic degradation pathways dominate at the other two sites.

In sediments of eutrophic, low-oxygen coastal systems, sulfate-dependent anaerobic oxidation of methane (S-AOM) is expected to account for most methane removal (Knittel and Boetius, 2009). Sulfate is presumably also the major terminal electron acceptor for AOM by ANME-2 archaea in the investigated sediments of the Stockholm Archipelago at all three sites. Estimated diffusive fluxes of sulfate and methane into the SMTZ (Table 3) suggest that S-AOM accounts for at least 40% of the observed sulfate reduction. Given the potential degassing of methane, especially for sediment intervals with high methane concentrations (Egger et al., 2016), *in situ* S-AOM rates are likely higher. The distinct gradient in calculated benthic effluxes of methane for our three sites suggest that the SMTZ loses its effectiveness as a filter for methane at the most sulfide-rich site (Table 3).

In coastal systems, electron acceptors other than sulfate may drive AOM, such as nitrate and nitrite (Haroon et al., 2013; Raghoebarsing et al., 2006) and easily reducible, poorly ordered Fe- and Mn-oxides (Aromokeye et al., 2020; Beal, House, & Orphan, 2009; Egger et al., 2015). At our sites, nitrate and nitrite are exclusively present in low concentrations in the surface sediment (Fig. 2) and are therefore unlike to substantially contribute to AOM activity. At Sites 5 and 7, easily reducible, poorly ordered Fe- and Mn-oxides are nearly absent (Fig. 3). Additionally, at both these sites, most of the reactive Fe and Mn is sulfidized, in line with the ambient bottom water redox conditions and relatively high porewater sulfide concentrations (Table 2; Fig. 2). Hence, there is only limited potential for Fe- and Mn-AOM, which seem to play a larger role in oligotrophic rather than eutrophic coastal ecosystems (Egger et al., 2015; Wytze K. Lenstra et al., 2018). However, a role for Fe-AOM cannot be fully excluded, as crystalline Fe-oxides (CDB Fe; Fig. 3) may also play a role in Fe-AOM (Bar-Or et al., 2017) and S-AOM (Sivan, Antler, Turchyn, Marlow, & Orphan, 2014). Magnetite, for example, was shown to stimulate Fe-AOM activity and ANME-2a enrichment in incubations with North Sea sediments (Aromokeye et al., 2020), and goethite-dependent AOM has been suggested as a significant methane sink in paddy soils, in which hematite and magnetite-AOM were also detected (He et al., 2021). Based on 16S rRNA gene profiling metal-AOM could occur, for example, below the SMTZ and the sulfide peak at Site 5, within the depth interval of 50-60 cm. Additionally, 16S rRNA gene profiling indicated that ANME-2 might persist below 50 cm at Site 7 (Fig. 4). At Site 3, some easily reducible, poorly crystalline Fe-oxides and Mn-oxides are present. Here, below 30 cm, ANME-2 16S rRNA gene-based relative abundances appear to increase concomitantly with the depletion of sulfate (Figure 5) and the appearance of the Fe^2+^ peak (Fig. 2). At Site 3, 16S rRNA gene-based and normalized genome coverage-based ANME-2 abundances seem to differ, likely due to shifts in the relative abundance of other microbial groups (Fig. 4), but general trends with depth are in agreement. The quantatitive importance of metal-AOM remains small, however, given the very low methane concentrations at Site 3.

Putative methanogen abundances (inferred from MAG coverage and 16S rRNA gene-based relative abundances) and methane production rates have contrasting profiles and do not positively correlate (Fig. 4). A potential explanation for these results is that larger pools of labile organic substrates generated from organic matter degradation could be available in surface sediments, which are depleted at depth, resulting in decreasing methane production rates in deeper sediment layers. In Site 3, where sulfate was detected until ca. 30 cm, sulfate reduction-driven thermodynamic inhibition of methanogenesis was expected (Bethke, Sanford, Kirk, Jin, & Flynn, 2011), and low methane production rates at this site indicate that this expectation was fulfilled despite the relatively high putative abundance of methanogens. In Sites 5 and 7, high methane production rates in surface sediments might also reflect larger pools of labile organic carbon and the rapid depletion of sulfate. These are conditions that could favor methanogens at potentially lower abundances in top sediments to be more active than in deeper sediments, where they could be more abundant but have less substrate availability. Additionally, the observation that the highest potential methane production rates were measured in surface sediments of Sites 5 and 7, concomitant with relatively high sulfate concentrations, suggests cryptic methane cycling. This could be fueled by methanogenesis from non-competitive substrates, a hypothesis that is supported by our data, since genomic potential for methanol and methylamine-driven methanogenesis was identified in three out of four methanogen MAGs. Methylotrophic methanogenesis has been previously implicated in cryptic methane cycling in similar ecosystems (Krause & Treude, 2021; Xiao et al., 2018).

Our key microbiological observation is the distribution of potential ANME-2 abundances based on MAG 11 normalized genome coverages. Low abundances of this organism were inferred in sediments of Site 3, in agreement with the low methane concentrations and potential methane production rates measured at this site (Fig. 4). At Site 5, ANME-2 abundances were the highest, within and below the SMTZ, matching abundant methane and sulfate substrates (Fig. 4) and their calculated fluxes into the SMTZ (Table 3), as well as highest potential methane production rates. However, at Site 7, where methane and sulfate were also abundant and had similar high fluxes into the SMTZ (Table 3), ANME-2 abundances were near zero. Based on our results, we infer that the distribution of ANME-2 at our study sites is most likely linked to high sulfide concentrations and a longer exposure time to sulfide at Site 7 (where concentrations were generally 0.5-1.2 mmol L^-1^), which could have directly caused sulfide toxicity in ANME cells. Additionally, sulfide could hamper the enzymatic activity of the F_420_-dependent sulfite reductase via product inhibition, leading to sulfite toxicity as well. This enzyme, described in *Methanocaldococcus jannaschii*, detoxifies sulfite by reducing it to sulfide, preventing intracellular reaction with proteins and sulfhydryl groups (Johnson & Mukhopadhyay, 2005). The disruption of the methane biofilter in the SMTZ likely explains the near 10-fold increase in the calculated benthic methane release from Site 5 to Site 7, 0.15 and 1.02 mmol m^-2^ day^-1^, respectively. Such high benthic fluxes of methane are in accordance with previously reported values for the Stockholm Archipelago (Sawicka & Brüchert, 2017).

MAG 11 ANME-2 had several genes encoding multiheme *c*-type cytochromes (Figure 3) including S-layer proteins, lowly expressed in Fe-AOM-performing ANME-2d (Cai et al., 2018), and OmcZ-like proteins recently suggested as the ANME mechanism for extracellular electron transfer (Chadwick et al., 2022), which could be used for electron transfer to a sulfate-reducing partner or to metal oxides. Multiheme *c*-type cytochromes identified in ANME might be targets of sulfite toxicity (Liu, Beer, & Whitman, 2012). Previous investigations have highlighted the role of Fe-AOM in coastal sediments (Aromokeye et al., 2020; Rooze, Egger, Tsandev, & Slomp, 2016), particularly in eutrophic areas where the SMTZ becomes shallower and thinner (Wallenius et al., 2021), as observed in this study (Figures. 2 and 4). F_420_-dependent sulfite reductases have been found to be highly abundant sulfur metabolism proteins in an ANME-2 metaproteomics study (Yu et al., 2018), which also reported the inhibition of AOM activity by ANME-2a/2c in methane seep sediments incubated with 1 mmol L^-1^ sulfite, 1 mmol L^-1^ polythionate and 0.25 mmol L^-1^ polysulfide. The authors concluded that the role of F_420_-dependent sulfite reductases in ANME-2 is more likely sulfite detoxification rather than sulfur assimilation, which could occur via several other ANME-2 enzymes (Yu et al., 2018). Furthermore, while methanogens are known to withstand 3-5 mmol L^-1^ sulfide levels and assimilate sulfur from sulfide (Liu, Beer, & Whitman, 2012), ANME archaea have been reported to be inhibited by 3-4 mmol L^-1^ sulfide under the low sulfate concentrations that we measured in our study (4 mmol L^-1^ range), but not under high (21 mmol L^-1^) sulfate (Timmers, Widjaja-Greefkes, Ramiro-Garcia, Plugge, & Stams, 2015). These previous studies suggest that a threshold sulfide concentration, potentially dependent on sulfate concentrations, might exert thermodynamic and toxicity controls on AOM activity. Our results match these previous observations, but also indicate that sulfide exposure time, which was more contrasting between Sites 5 and 7 than sulfide concentrations, could play a role in the lower ANME-2 abundances at Site 7 (Fig. 4). Overall, these results support our biogeochemical and metagenomic evidence for the proposed mechanism of sulfide toxicity as a key control on ANME-2 abundances and activity in coastal sediments, suggesting that the expansion of euxinia in coastal areas (Breitburg et al., 2018) might increase benthic methane release into the water column and potentially coastal methane emissions to the atmosphere. This is particularly relevant for relatively shallow coastal sites, such as Site 7 in the Stockholm Archipelago. Genes encoding F_420_-dependent sulfite reductases have been found in ANME-1, ANME-2 and ANME-3 genomes (Chadwick et al., 2022), suggesting that sulfide-driven sulfite toxicity may be a commonly encountered environmental pressure by ANME and, therefore, a widespread control on AOM activity.

## Conclusions

Our results indicate that bottom water hypoxia and euxinia at eutrophic sites in the Stockholm Archipelago induced shifts in sediment geochemistry, microbial community structure, and methanogenic activity, resulting in increased potential methane production rates and *in situ* methane concentrations. Our data suggest that ANME-2 archaea might be able to compensate for methane increases under hypoxic conditions, but are unable to thrive under euxinic conditions because of sulfide toxicity. This disruption of the methane biofilter may result in increased benthic methane release into the water column in coastal ecosystems severely impacted by eutrophication and bottom water deoxygenation. Further studies should investigate if increased methane concentrations in euxinic waters result in increased methane emissions to the atmosphere. Moreover, future studies are required to characterize the methane-oxidizing activity of ANME archaea under changing bottom water redox conditions, as well as the metabolism and terminal electron acceptors utilized.

## Acknowledgements

We thank the captain, the crew of RV Electra and Christoph Humborg for their assistance with the fieldwork. Arnold van Dijk, Coen Mulder, Helen de Waard, Tom Bastiaan and Liam Kirwan (Utrecht University) are thanked for analytical assistance. This study was funded by ERC Synergy MARIX 854088 (MSMJ, CPS and NAGMvH), NESCC 02001001 (WKL, AJW, MSMJ & CPS), SIAM 024002002 (MJEM, CUW, MSMJ), Havsochvattenmyndigheten DNR 1960-2018 (MH & CPS), and VI.Veni.212.040 (PDM). We are also grateful to Rasmus Rodineliussen for assistance with sampling, Linnea Kop for ideas and code for data visualization, and Mike Lee for creating the DADA2 tutorial (https://astrobiomike.github.io/amplicon/dada2_workflow_ex) from which we derived code for the 16S rRNA gene analyses in this study.

## List of Supplemental Materials

**Supplemental Table 1**. Geochemical data.

**Supplemental Table 2**. List of MAG 011 ANME-2 genes and corresponding loci identified in this study.

**Supplemental Table 3**. Calculations of methane production rates in sediment incubations.

**Supplementary Figure 1.**
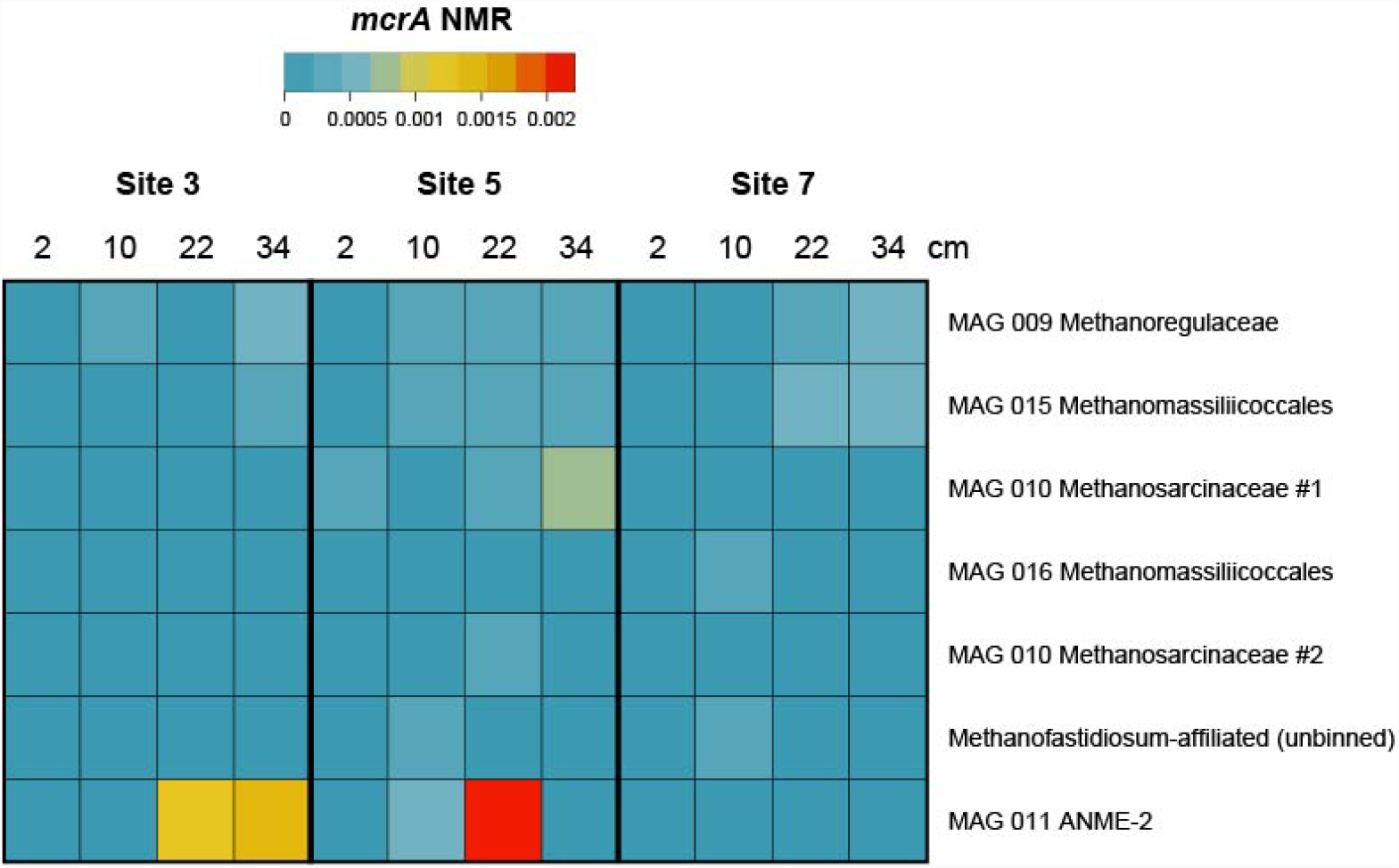
Heat map of *mcrA* NMR values across sites and depths.

**Supplementary Figure 2.**
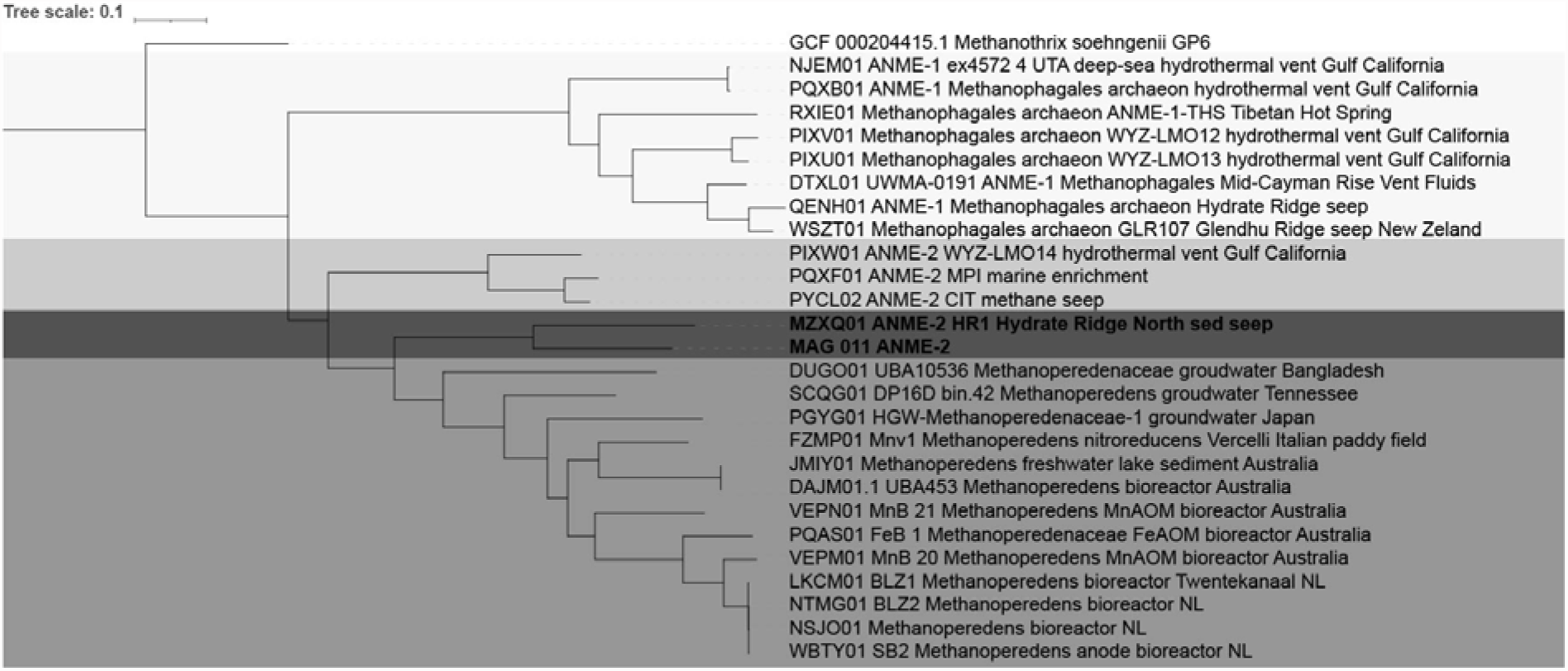
UBCG tree with 92 concatenated archaeal genes of reference ANME genomes (as indicated by NCBI accession numbers) and the ANME-2 genome from this study. No bootstrap values were larger than 0.7.

**Supplementary Figure 3.**
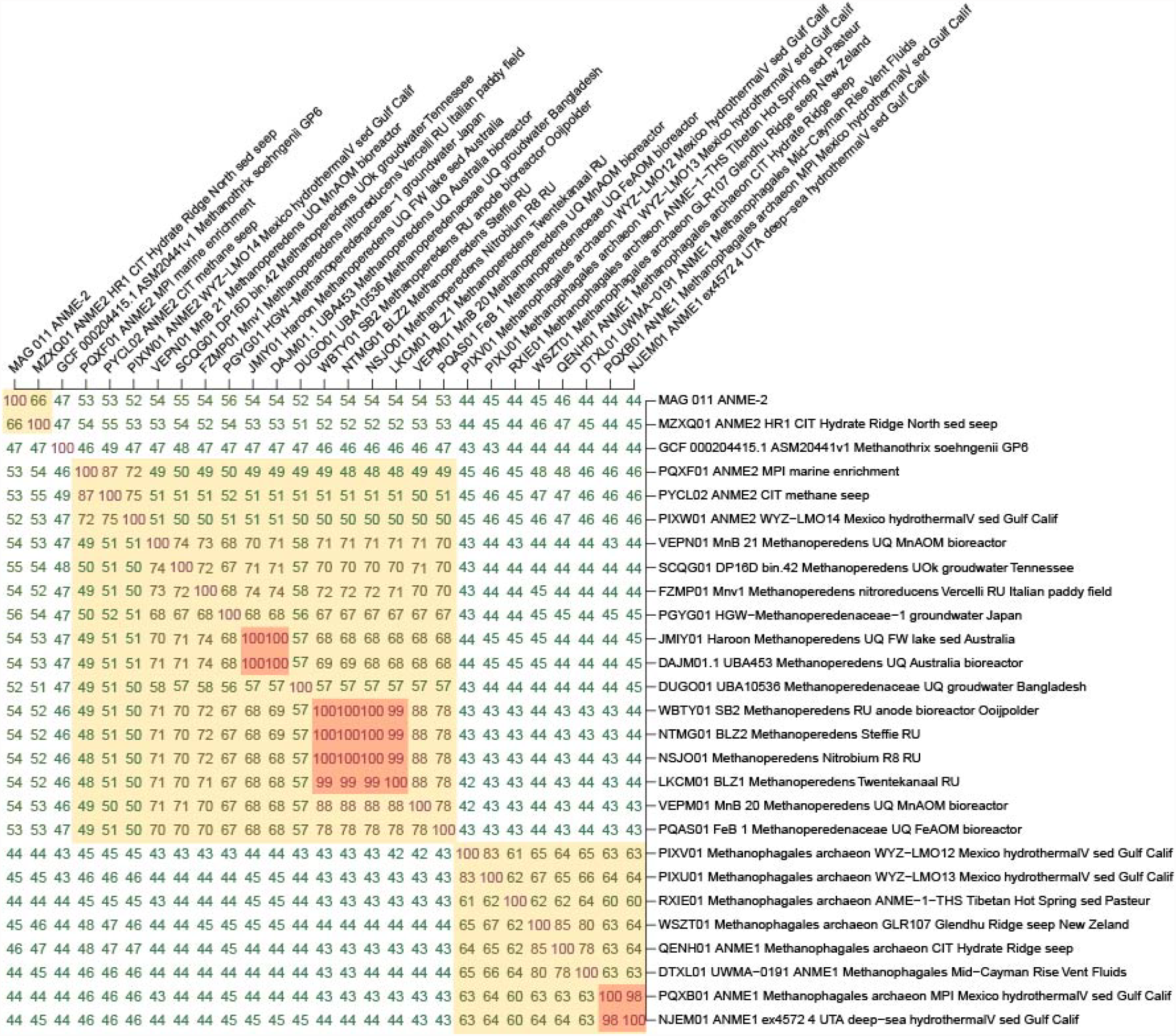
Average amino acid identity matrix with pairwise comparisons among genomes presented in Supplemental Figure 2.

